# Efficient reaction deletion algorithms for redesign of constraint-based metabolic networks for metabolite production with weak coupling

**DOI:** 10.1101/563437

**Authors:** Takeyuki Tamura

## Abstract

**Background:** Metabolic network analysis through flux balance is an established method for the computational redesign of production strains in metabolic engineering. The computational redesign is often based on reaction deletions from the original wild type networks. A key principle often used in this method is the production of target metabolites as by-products of cell growth. From a viewpoint of bioinformatics, it is very important to prepare a set of algorithms that can determine reaction deletions that achieve growth coupling whatever network topologies, target metabolites and parameter values will be considered in the future. Recently, the strong coupling-based method was used to demonstrate that the coupling of growth and production is possible for nearly all metabolites through reaction deletions in genome-scale metabolic models of *Escherichia coli* and *Saccharomyces cerevisiae* under aerobic conditions. However, when growing *S. cerevisiae* under anaerobic conditions, deletion strategies using the strong coupling-based method were possible for only 3.9% of all metabolites. Therefore, it is necessary to develop algorithms that can achieve growth coupling by reaction deletions for the conditions that the strong coupling-based method was not efficient.

**Results:** We developed an algorithm that could calculate the reaction deletions that achieve the coupling of growth and production for 91.3% metabolites in genome-scale models of *S. cerevisiae* under anaerobic conditions. This analysis was conducted for the worst-case-scenario using flux variability analysis. To demonstrate the feasibility of the coupling, we derived appropriate reaction deletions using the new algorithm for target production in which the search space was divided into small cubes (CubeProd).

**Conclusions:** We developed a novel algorithm, CubeProd, to demonstrate that growth coupling is possible for most metabolites in *S.cerevisiae* under anaerobic conditions. This may imply that growth coupling is possible by reaction deletions for most target metabolites in any genome-scale constraint-based metabolic networks. The developed software, CubeProd, implemented in MATLAB, and the obtained reaction deletion strategies are freely available.

## 1 Background

Computational models are gaining importance in the metabolic engineering field. These models can design and optimize metabolite producing microbes [1–3]. Coupling cellular growth with the production of a desired metabolite is a key principle, known as growth coupling, in computational strain designs [4]. Unfortunately, growth coupling is not possible for every metabolite.

Flux balance analysis (**FBA**) is a widely used mathematical model in metabolic engineering simulations. FBA assumes a pseudo-steady state in which the sum of incoming fluxes must equal the sum of outgoing fluxes for each internal metabolite [5]. FBA can be formalized as linear programming (**LP**). In the LP, biomass production flux is maximized. The obtained maximum value is referred to as the growth rate (**GR**). For each metabolite, the production rate (**PR**) is estimated based on the assumption that GR is maximized. When both the GR and PR exceed certain thresholds, growth coupling is achieved.

Many efficient solvers are available for LP, as LP is polynomial-time solvable. Therefore, FBA simulations are efficiently possible even for genome-scale metabolic models. The fluxes calculated by FBA correspond to experimentally obtained fluxes [6].

With FBA, numerous computational methods can be used to identify optimal design strategies including reaction deletions. OptKnock is a well-established method that identifies optimal design strategies [7]. OptKnock derives its design strategies through bi-level linear optimization using mixed integer LP. However, because mixed integer LP is an NP-complete problem [8], OptKnock cannot identify design strategies for many genome-scale networks. Many methods have been developed to overcome this problem. OptGene and Genetic Design through Local Search (GDLS) identify gene deletion strategies using a genetic algorithm and a local search with multiple search paths, respectively [9, 10]. EMILio and Redirector use iterative linear programs [11, 12]. Genetic Design through Branch and Bound (GDBB) uses a truncated branch and branch algorithm for bi-level optimization [13]. Fast algorithm of knockout screening for target production based on shadow price analysis (FastPros) is an iterative screening approach that discovers reaction knockout strategies [14]. IdealKnock utilizes the ideal-type flux distribution and ideal point=(GR, PR) [15]. Parsimonious enzyme usage FBA (pFBA) [16] finds a subset of genes and proteins that contribute to the most efficient metabolic network topology under the given growth conditions. GridProd integrates the ideas of IdealKnock and pFBA [17].

In 2017, von Kamp and Klamt developed a strong coupling-based method and showed that growth coupling is possible for more than 90% of all target metabolites by reaction deletions in *Escherichia coli* and *Saccharomyces cerevisiae* under aerobic conditions. Similar results were provided also for *Aspergillu niger, Corynebacterium glutamicum*, and *Synechocystis*. They also showed that growth coupling is possible for more than 75% of all target metabolites in *E. coli* under anaerobic conditions. However, for *S. cerevisiae* under anaerobic conditions, the strong coupling-based method could only derive reaction deletion strategies for 3.9% of all metabolites. They also showed that strong coupling is impossible for any gene-associated reaction deletion strategy that achieves a target production exceeding 10% of the theoretical maximum yield (**TMY**) for 94.6% of all metabolites. This does not mean that growth coupling is impossible for 94.6% of all target metabolites, but strong coupling by gene-associated reaction deletions is impossible. When strong coupling is achieved, PR exceeds certain criteria for any flux even when GR is not maximized [18]. Therefore, growth coupling that is not strong coupling may be possible particularly if any reaction deletion is allowed. Due to development of chemical engineering, deletion of any reaction may become possible in near future. In this study, we developed a novel algorithm, CubeProd, for finding reaction deletion strategies to demonstrate that growth coupling is possible for most metabolites in *S.cerevisiae* under anaerobic conditions.

## 2 Results

The associated FBA model for *S. cerevisiae* was iMM904, that contains 1228 internal metabolites. However, the TMY is zero for 541 of the 1228 metabolites. Therefore, we applied CubeProd for the 687 remaining target metabolites in iMM904 which were the same as those evaluated by von Kamp and Klamt [4]. The model configurations in the computational experiments were the same as in [4], but any reaction deletion strategy was allowed.

Computations were conducted for three different levels of demanded minimum product yield, 10%, 30%, and 50% of the TMY for each target metabolite to compare with the results of [4]. Additionally, the experiments for 1% of the TMY were also conducted to investigate the performance of CubeProd. When the strong coupling-based method was applied, suitable reaction deletion strategies were obtained for only 3.9%, 1.7% and 1.3% of target metabolites for the 10%, 30%, and 50% minimum yield levels, respectively [4]. However, when CubeProd was applied, suitable reaction deletion strategies were found for 91.3%, 69.6%, and 47.6% of all target metabolites for the 10%, 30% and 50% minimum yield levels, respectively, as shown in Fig. 1. While the numbers of the reaction deletions derived through the strong coupling-based method were less than 3% of all reactions, those derived through CubeProd were 78.9% on average.

**Figure 1.**
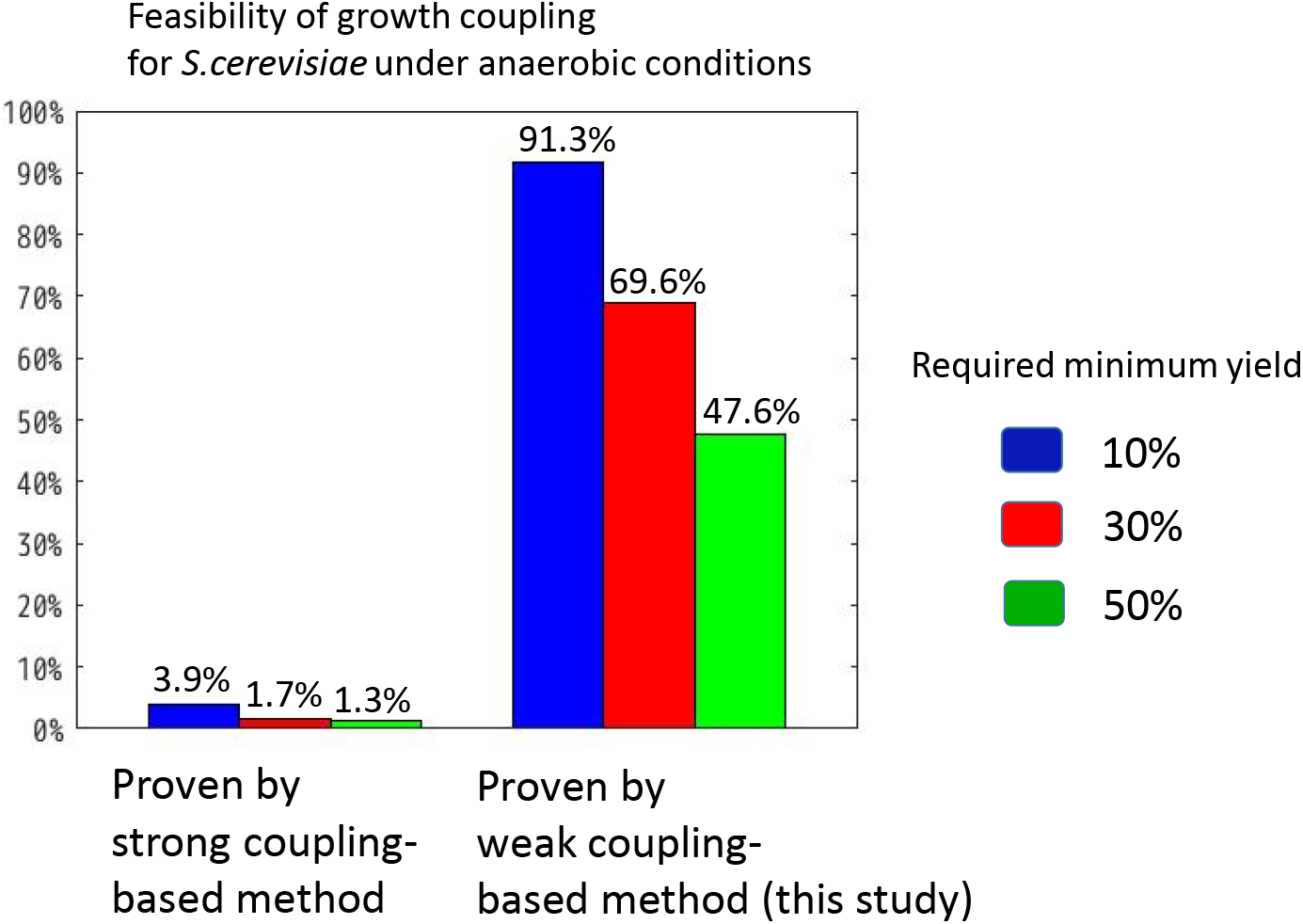
Feasibility of growth coupling for *S. cerevisiae* under anaerobic conditions demonstrated by the strong coupling-based method and CubeProd. The feasibility was demonstrated by calculating reaction deletion strategies. Percentage of producible organic metabolites is shown for each required minimum yield.

All procedures for CubeProd were implemented on a CentOS 7 machine with an Intel Xeon Processor with 2.30 GHz 18C/36T, and 128 GB memory. This workstation had CPLEX, COBRA Toolbox [19], and MATLAB.

Fig. 2 shows the percentages of producible organic metabolites by the reaction deletion strategies obtained by CubeProd that achieved growth coupling for the 687 target metabolites of iMM904 for different cube sizes and demanded minimum yield levels. The horizontal axis corresponds to different resolutions (*P*^−1^) of CubeProd. The blue, red, black, and green lines correspond to the demanded minimum yield levels of 1%, 10%, 30%, and 50%, respectively. When the demanded minimum yield was 1% of the TMY, the percentage was 88.9% with *P*^−1^ = 5 and 93.2% with *P*^−1^ ≤ 15. When the demanded minimum yield was 10% of the TMY, the percentage was 76.0% with *P*^−1^ = 5, 90.8% with *P*^−1^ ≤ 15, and 91.4% with *P*^−1^ ≤ 30. When the demanded minimum yield was 30% of the TMY, the percentage was 51.1% with *P*^−1^ = 5 and 76.6% with *P*^−1^ ≤ 15. When the demanded minimum yield was 50% of the TMY, the percentage was 25.0% with *P*^−1^ = 5 and 59.7% with *P*^−1^ ≤ 15. Each line is shown with a monotone increase because the experiments were conducted for every 5 ≤ *c* ≤ *P*^−1^ when *P*^−1^ = *c*. The average computation time for each producible metabolite was 39.7s and 943.7s for *P*^−1^ = 5 and *P*^−1^ = 15, respectively.

**Figure 2.**
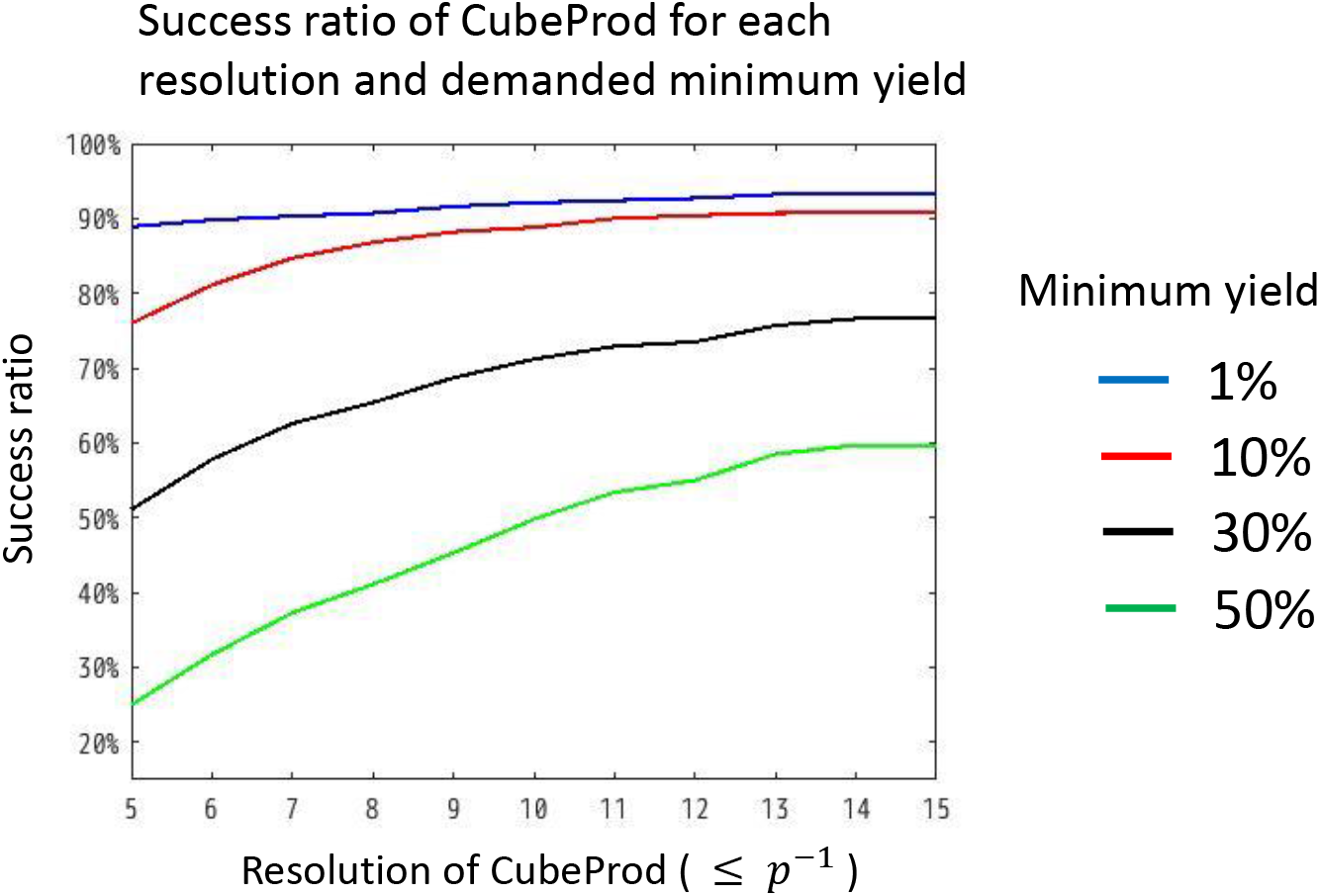
Success ratio of CubeProd in identifying reaction deletion strategies that could achieve growth coupling for 687 target metabolites of iMM904 at each resolution (≤ *P*^−1^) and demanded minimum yield.

## 3 Discussion

Fig. 1 shows that CubeProd identified reaction deletion strategies that can achieve growth coupling for 91.3%, 69.6%, and 47.6% of the 687 target metabolites in iMM904 for the demanded minimum yield levels of 10%, 30%, and 50%, respectively. The strong coupling-based method could identify reaction deletion strategies for 3.9%, 1.7%, and 1.3% of metabolites, respectively [4]. Because the strategies by CubeProd include non-gene associated reactions and the average deletion size of CubeProd is 78.9%, readers may think that the strategies by CubeProd are not realistic. However, whether each reaction is gene-associated is according to current knowledge and technologies, which may be updated in future. Therefore, it is important to prove the possibility of growth coupling when any reaction deletion is allowed. Computational experiments using CubeProd showed that growth coupling is possible for most metabolites, even in *S. cerevisiae* under anaerobic conditions.

Strong coupling was defined in [18]. Biomass synthesis is strongly coupled with product synthesis if any steady-state flux has a product yield greater than or equal to the demanded minimum yield and at least one steady-state flux has a biomass yield greater than or equal to the demanded minimum growth rate. This study also defined weak coupling. Biomass synthesis is weakly coupled with product synthesis if any steady-state flux with a biomass yield greater than or equal to the demanded minimum growth rate exhibits a product yield of greater than equal to the demanded minimum product yield and at least one such flux exists.

Growth coupling is defined under the condition that GR is maximized. If the demanded minimum growth rate equals the maximum growth rate, the definition of weak coupling becomes the same as that of growth coupling. In this study, the reaction deletion strategies that achieve growth coupling were determined. In other words, the reaction deletion strategies that achieve a special case of weak coupling were determined.

### 3.1 Structural properties

von Kamp and Klamt explained what structural properties of the yeast metabolism resulted in impossibility of strong coupling by gene-associated reaction deletions as follows [4]. Because excretion of ethanol is essential for anaerobic growth on glucose [20], the reaction steps from glucose to ethanol are essential and must be kept active. Therefore, no suitable deletion sets can then exist that would guarantee a minimum yield of the respective metabolite because the substrate glucose, in principle, be completely converted to ethanol.

However, this is not an issue for deletion strategies including non-gene-associated reactions. Computational experiments were conducted to investigate the activity of the ethanol exchange reaction by deleting the reactions described in Fig. 3(A), which shows the network topology around the exchange reaction of ethanol in iMM904. “EX_etoh_e” represents the reaction of “ethanol exchange” that is an irreversible excretion reaction. The substrate of “EX_etoh_e’ is “etoh_e.” “ETOHt” is the ethanol reversible transport reaction between “etho_e” and “etoh_c” Both “etoh_e” and “etoh_c” represent ethanol. The pathway from “EX_etoh_e” to “etoh_c” is in the chain structure. However, “etoh_c” has seven adjacent reactions “ETOHt,” “ACHLE3,” “ALCD2x_copy1,” “ALCD2ir,” “ALCD2x_copy2,” “ETOHtm,” and “OHACT1” that represent ethyl acetate hydrolyzing esterase, alcohol dehydrogenase (ethanol), alcohol dehydrogenase reverse rxn acetaldehyde ethanol, alcohol dehydrogenase (ethanol), ethanol transport to mitochondria diffusion, alcohol acetyltransferase ethanol, respectively. “ACHLE3” and “ALCD2ir” are irreversible reactions to “etoh_c.” “ALCD2x_copy1” and “OHACT1” are irreversible reactions from “etoh_c.” “ALCD2x_copy2” and “ETOHtm” are reversible reactions.

**Figure 3.**
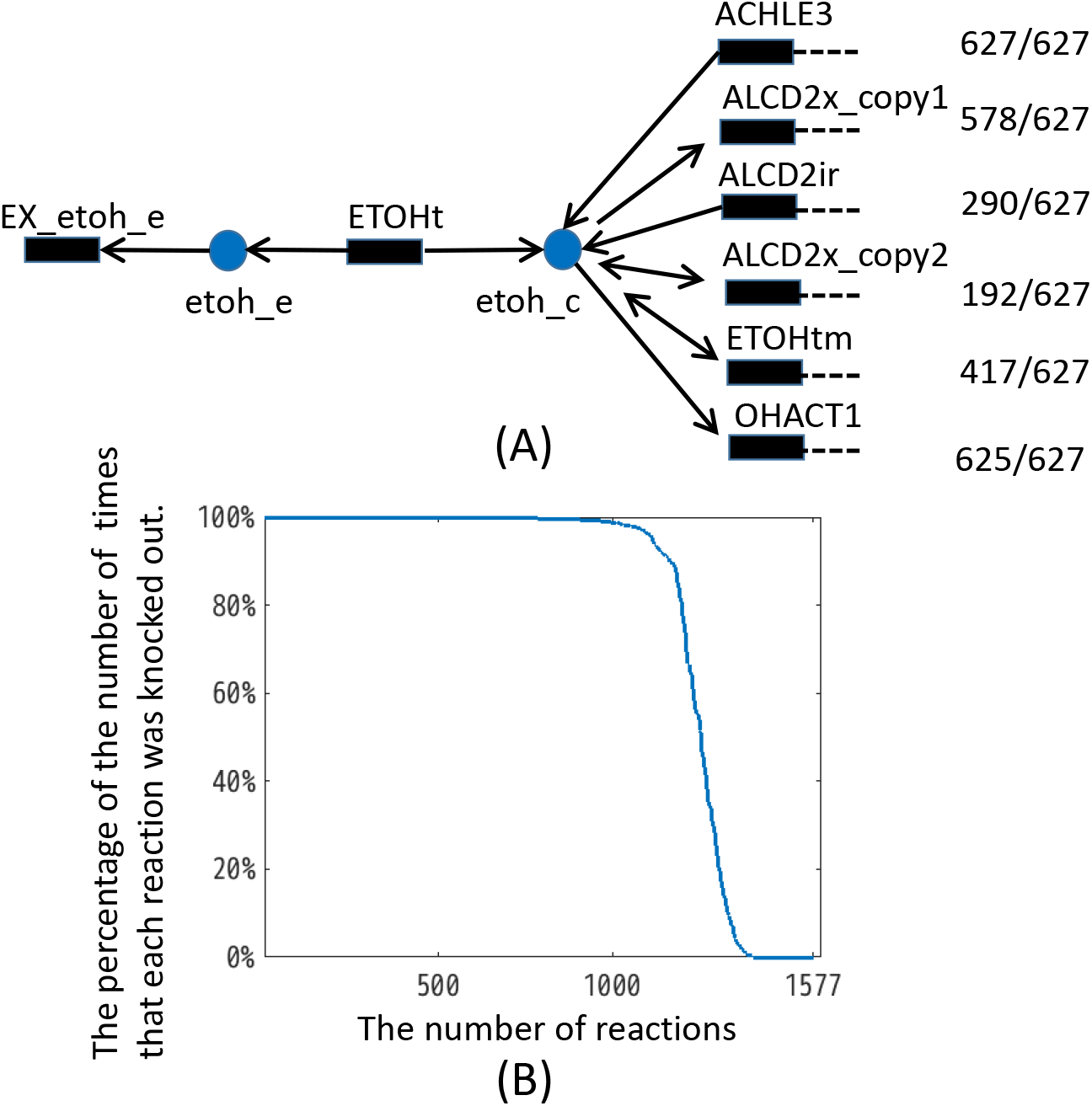
(A) Network topology of iMM904 around ethanol exchange reaction, and the number of times that the six reactions excluding “EX_etho_e” and “ETOHt” were deleted in the CubeProd strategy. (B) The percentage of the number of times each reaction was deleted in the CubeProd strategy.

In wild type, when GR was maximized, the GR of iMM904 in anaerobic condition was 0.2879, and the minimum value of the ethanol exchange was 15.81. When all the six reactions in Fig. 3(A) excluding “EX_etoh_e” and “ETOHt” were deleted, GR became 0.2311 and the maximum and minimum values of ethanol exchange were 0. Therefore, for iMM904 in anaerobic conditions, the excretion of ethanol exchange was not essential when GR is maximized.

Next, the cases where five of the six reactions were deleted were simulated in iMM904. The minimum and maximum amounts of ethanol exchange evaluated by FVA became 0 for three of the six cases, “only ACHLE3 is active,” “only ALCD2x_copy1 is active,” and “only OHACT1 is active.” However, the amount of the ethanol exchange was the same as that of the wild type for the other three cases, “only ACLD2ir is active,” “only ALCD2x_copy2 is active,” and “only ETOHtm is active.” For the 627 target metabolites for which CubeProd found the deletion strategies, “ACHLE3,” “ALCD2x_copy1,” “ALCD2ir,” “ALCD2x_copy2,” “ETOHtm,” and “OHACT1” were deleted in 627, 578, 290, 192, 417, and 625 cases, respectively. Therefore, it is possible to delete reactions between glucose and ethanol to achieve growth coupling that is not strong coupling in iMM904 under anaerobic conditions.

### 3.2 Size of deletion

Fig. 3(B) represents the number of times each reaction was deleted when CubeProd succeeded in finding deletion strategies. Five hundred and forty-eight of the 1577 iMM904 reactions were deleted for all 627 cases where CubeProd found appropriate deletion strategies. One hundred and seventy-three of the 1577 iMM904 reactions were never deleted. On average, 1245 of the 1577 reactions (78.9%) were deleted for each target metabolite.

Finding knockout strategies with minimum sets of genes or reactions to produce valuable metabolites has been an important problem in computational biology. Because large amounts of time and effort are required to experimentally knock out several genes, a smaller number of knockouts is preferred. Fortunately, DNA synthesis technologies are consistently being improved [21]. Designing synthetic short DNA may replace the method of knocking out genes from the original long genome to produce metabolites. In such a case, the number of genes included in the design of synthetic DNA should be as small as possible because of the experimental effort and time required.

### 3.3 Comparing CubeProd with the strong coupling-based method and GridProd

Identifying an optimal subnetwork that achieves the maximum PR when GR is maximized is an NP-hard problem; thus, it is often impossible to determine this information for genome-scale models in a realistic time. To overcome this problem, the author previously developed GridProd, which does not guarantee the identification of the optimal subnetwork, but can reveal appropriate deletion strategies for many target metabolites for *E.coli* under microaerobic conditions [17]. However, GridProd was not efficient for *S.cerevisiae* under anaerobic conditions in preliminary experiments.

When GridProd could not derive appropriate reaction deletion strategies, the fluxes obtained by the LP were often quite different from those obtained by the FVA. This is because the optimal solution of LP is not always uniquely determined. One of the most promising methods for overcoming this problem is to decrease the number of optimal solutions by adding other constraints for each LP.

To this end, while GridProd imposes the following two constraints

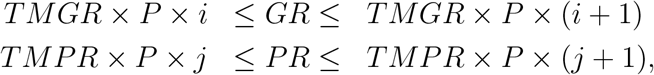

CubeProd imposes the following three constraints

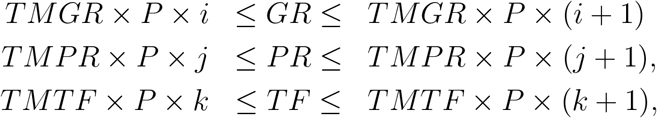

where TMTF represents the theoretical maximum total flux for all integers 1 ≤ *i,j,k* ≤ *P*^−1^. CubeProd minimizes the total amount of absolute amounts of fluxes.

The concept of CubeProd can be compared to the strong coupling-based method and GridProd using the following examples on the toy model of the metabolic network shown in Fig. 4. {R1,…,R8} and {C1,…,C5} are sets of reactions and metabolites, respectively. All internal reactions are gene-associated. R1 is a source exchange reaction such as glucose or oxygen uptake. R2 consumes C1 and produces the same amount of C2. R3 is the biomass objective function consuming C2 that corresponds to GR. R4 consumes C1 and produces the same amount of C3. R5 consumes C3 and produces the same amount of C4. R6 consumes C4 and produces the same amounts of C2 and C5. R7 consumes C1 and produces the same amount of C5. R8 is the exchange reaction of the target metabolite C5. All of the reactions are irreversible. [*a, b*] indicates that *a* and *b* are the lower and upper bounds of the flux for the corresponding reaction, respectively. The minimum required GR and PR are two. R1, R3, and R8 are not allowed to be deleted.

**Figure 4.**
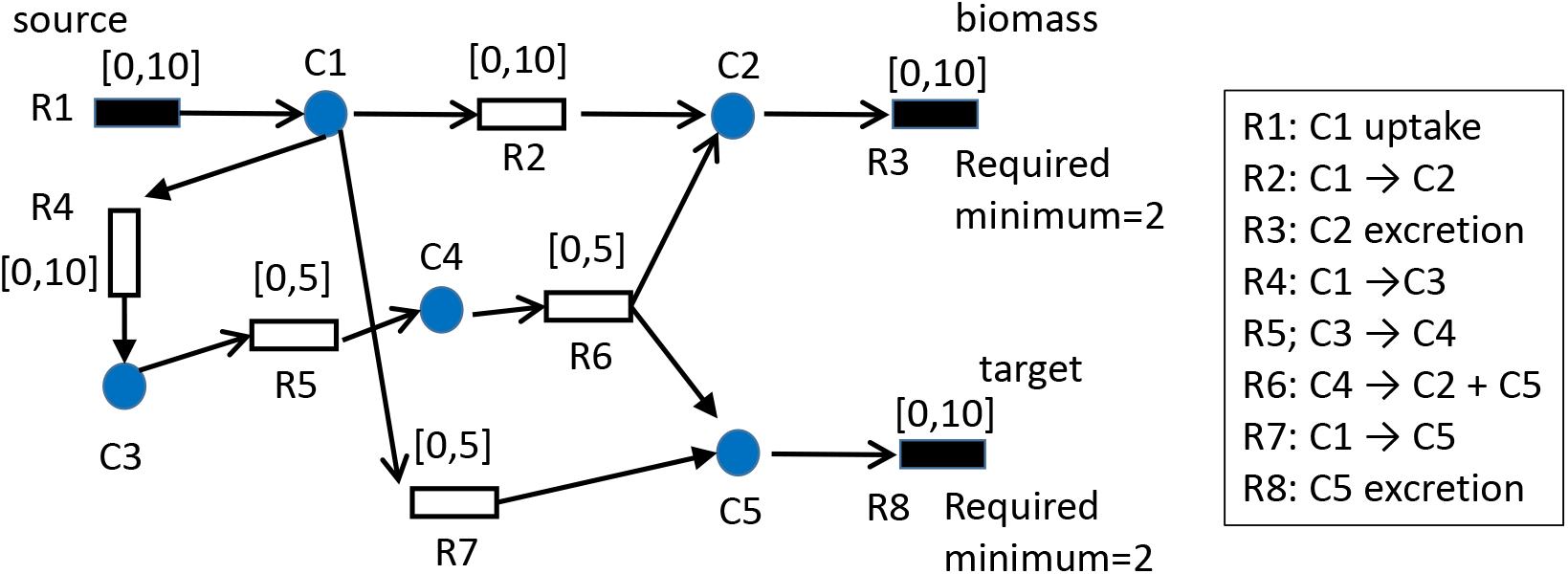
A toy example in which CubeProd can identify the optimal strategy but either the strong coupling-based method or GridProd cannot.

Under the assumption that GR is maximized, the optimal design strategy for the worst case-scenario by FVA is to “delete R2” or “delete R2 and R7.” When R2 is deleted, R3 becomes 5, and 5 is produced for C5 and R8 via R6. In the best case-scenario, another 5 is produced for C5 and R8 via R7. However, R7 becomes 0 in the worst case. Therefore, when only R2 is deleted, (GR, PR)=(5,5) is obtained by FVA for the worst case. Similarly, when R2 and R7 are deleted, (GR, PR)=(5,5) is obtained. When R2 is active and R3 is maximized, R3 becomes 10 and R8 becomes 0. This is because the upper bounds of R1, R2, and R3 are 10. When some of {R4, R5, R6} is deleted in addition to R2, GR=0 is obtained and the required minimum is not satisfied.

In the following, we investigate whether the strong coupling-based method, CubeProd, and GridProd can identify the optimal strategies in this example.

Strong coupling demands that the product yield is more than the required minimum for all non-zero flux vectors. When all R4, R5 and R6 are active, (R3,R8) = (1,1) is possible by (R2,R4,R5,R6,R7) = (0,1,1,1,0), but R8=1 does not satisfy the required minimum for PR. Therefore, strong coupling is impossible when all R4, R5 and R6 are active. Suppose that either R4, R5 or R6 is deleted. When R2 is deleted, it is impossible to satisfy the required minimum for GR. When R7 is deleted, (R3,R8)=(1,0) is possible, and does not satisfy the required minimum for either GR or PR. Thus, strong coupling is infeasible for the network of Fig. 4 for any reaction deletion. Therefore, the strong coupling-based method cannot find either the optimal strategy or strategy satisfying the required minimums.

GridProd predicts the design strategy for each small ranges of GR and PR. For example, suppose that 4 ≤ GR ≤ 6 and 4 ≤ PR ≤ 6 are considered. GridProd determines a flux vector satisfying 4 ≤ GR ≤ 6 and 4 ≤ PR ≤ 6 under the condition that the total amount of fluxes is minimized. For simplicity of explanation, the total amount of fluxes of only internal reactions is considered here. In this case, (R2,R4,R5,R6,R7)=(4,0,0,0,4) is determined. Because (R4,R5,R6) = (0,0,0), GridProd deletes R4, R5 and R6. This results in (GR,PR)=(10,0) under the condition that GR is maximized. Even when the resolution is infinity and GR=5 and PR=5 are considered, (R2,R4,R5,R6,R7)=(5,0,0,0,5) is determined and (GR,PR)=(10,0) is obtained. Because GridProd minimizes the total amount of fluxes, R4, R5, and R6 are preferentially deleted instead of R2. When R2 is active, (GR,PR)=(10,0) is always obtained under the condition that GR is maximized. Thus, GridProd cannot find either the optimal strategy or strategy satisfying the required minimums.

CubeProd predicts the design strategy for each small ranges of GR, PR, and the total amount of fluxes (TAF). For example, suppose that 5 − *ϵ* ≤ GR ≤ 5 + *ϵ*, 5 − *ϵ* ≤ PR ≤ 5 + *ϵ*, and 15 − *ϵ* ≤ TAF ≤ 15 + *ϵ* are considered, where *ϵ* is a small positive constant. For simplicity of explanation, the TAF of the internal reactions is considered and *t* is zero. Then, the flux must satisfy R2 + R5 =5, R5 + R7 = 5, R2 + 3·R5 + R7 =15, and (R2,R4,R5,R6,R7)=(0,0,5,0,0) is uniquely determined. Therefore, R2 and R7 are deleted, and (GR,PR)=(5,5) is obtained under the condition that GR is maximized.

Suppose that the resolution of CubeProd is not large enough. For example, suppose that 4 < GR ≤ 6,4 ≤ PR ≤ 6, 12 ≤ TAF ≤ 18 are considered. In this case, (R2,R4,R5,R6,R7) = (6,0,0,0,6) also satisfies the constraints in addition to (R2,R4,R5,R6,R7) = (0,4,4,4,0). If CubeProd selects the former, (GR,PR)=(10,0) is obtained, although (GR,PR)=(5,5) is obtained if CubeProd selects the latter.

Thus, for the network of Fig. 4, either the strong couple-based method or GridProd cannot find the appropriate strategies, but CubeProd can if the resolution is large enough.

## 4 Conclusion

We developed a novel algorithm, CubeProd, and demonstrated that growth coupling is possible for most metabolites in *S.cerevisiae* under anaerobic conditions, for which the strong coupling-based method was not efficient. The determined reaction deletion strategies by CubeProd include non-gene-associated reactions and gene-associated reactions. Current knowledge determines whether each reaction is gene-associated, and this knowledge may be updated or refined in future. Thus, it is also important to consider reaction deletion strategies including non-gene-associated reactions.

CubeProd determines reaction deletion strategies that achieve growth coupling for genome-scale metabolic networks by extending the concept of GridProd [17], which was developed by the author. GridProd divides the solution space of optKnock into *P*^−2^ small grids and conducts LP and flux variability analysis (**FVA**) for each grid, where *P*^−1^ is a positive integer parameter. The LP identifies the reactions to be deleted. The FVA calculates the minimum PR of the target metabolite when the determined reactions by the LP are deleted and GR is maximized for each grid. The deletion strategy of the grid whose PR is highest is then adopted as the GridProd solution. The area size of each grid in GridProd is (*P* × *TMGR*) × (*P* × *TMPR*), where TMGR and TMPR are the theoretical maximum growth rate and theoretical maximum production rate, respectively.

In addition to (*P* × *TMGR*) × (*P* × *TMPR*), CubeProd divides the solution space by (*P* × *TMGR*) × (*P* × *TMPR*) × (*P* × *TMTF*), where TMTF is the theoretical maximum total flux.

## 5 Methods

### 5.1 Genome-scale metabolic model of *S. cerevisiae*

We used iMM904 as an original mathematical model of metabolic networks [22]. iMM904 is a genome-scale reconstruction of the metabolic network in *S. cerevisiae*. This model includes 1228 internal metabolites and 1577 reactions with 1020 gene-associated reactions and 557 non-gene associated reactions. The upper bound of glucose uptake was 10 mmol gDW^−1^h^−1^, and the ATP maintenance demand was 1 mmol gDW^−1^h^−1^. To simulate anaerobic conditions, cytochrome *c* oxidase was removed as in [4]. To simulate the production potential of each target metabolite in this model, an exchange reaction was temporarily added to the model as in [4].

### 5.2 Pseudo-code of CubeProd

The pseudo-code of CubeProd is as follows.

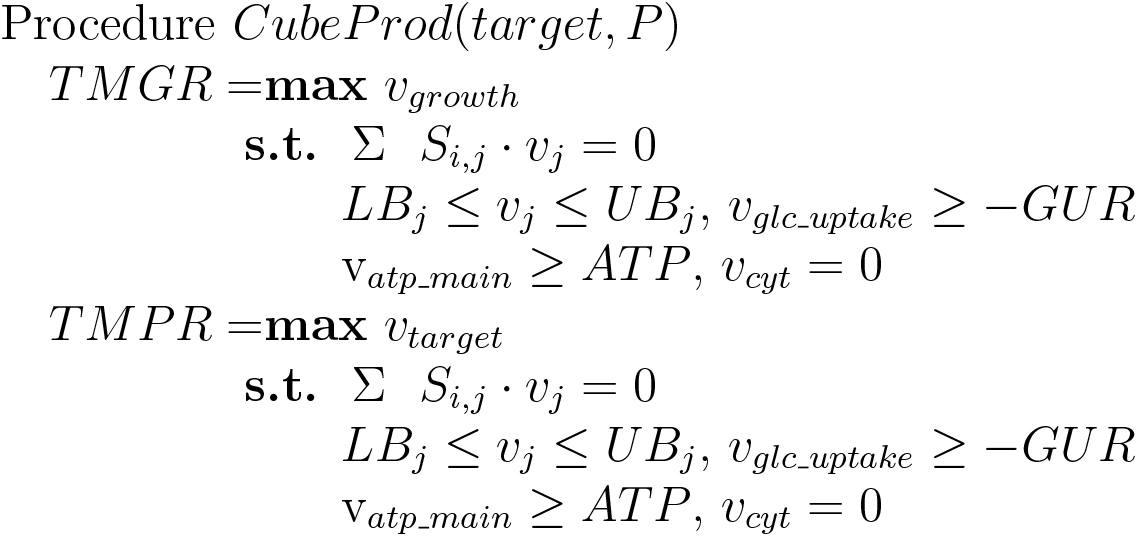
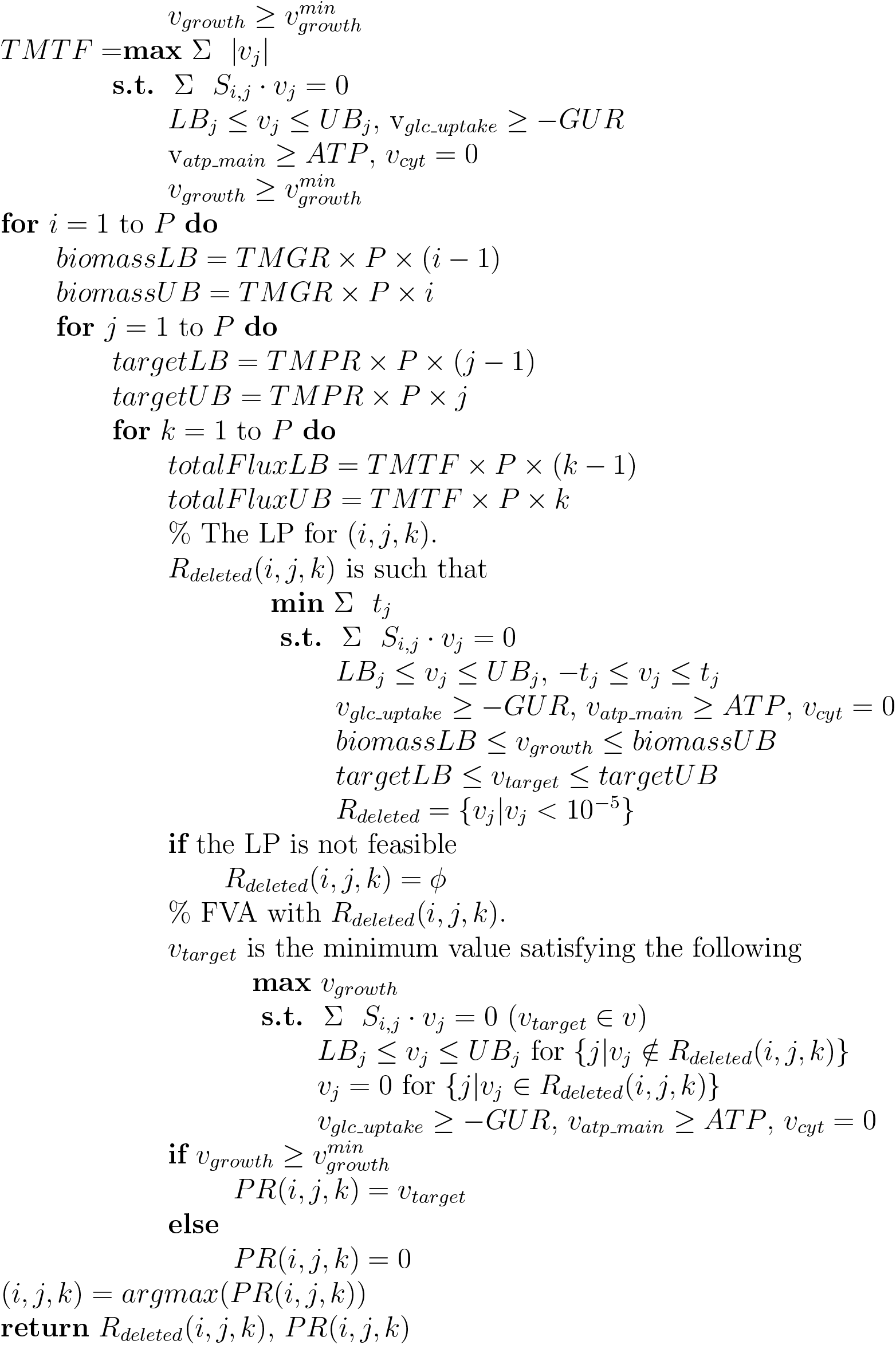

In the above pseudo-code, the TMGR, TMPR, and TMTF are calculated first. *S_i,j_* is the stoichiometric matrix. *LB_j_* and *UB_j_* are the lower and upper bounds of *v_j_*, respectively, which represent the flux of the *j*th reaction.

*v_glc_uptake_*, and *v_atp_main_* are the lower bounds of the uptake rate of glucose (GUR), and ATP maintenance demand, respectively. 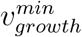 is the minimum required cell growth rate.

In each cube, LP and FVA are conducted. “*biomassLB*” and “*biomassUB*” represent the lower and upper bounds of GR, respectively. “*targetLB*” and “*targetUB*” represent the lower and upper bounds of PR, respectively. *“totalFluxLB”* and *“totalFluxUB”* represent the lower and upper bounds of total amounts of fluxes, respectively. These values are used as the constraints in the LP. Each cube is represented by the three constraints, “*biomassLB ≤ v_growth_ ≤ biomassUB*,” “*targetLB* ≤ *v_target_* ≤ *targetUB*,” and “*totalFluxLB* ≤ £ |*v_j_*| ≤ *totalFluxUB*.” *TMPR* × *P*, *TMGR* × *P*, and *TMTF* × *P*, represent the length of the first, second and third dimension for each cube, respectively.

In the solution of the LP, a set of reactions whose fluxes are nearly 0 (less than 10^−5^) are represented as *R_deleted_*, which is used as a set of deleted reactions in the FVA. In the FVA, the fluxes of the reactions included in *R_deleted_* were forced to be 0. If the obtained PR is more than or equal to 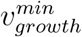 in the solution of the FVA, the value of PR is stored to *PR*(*i, j, k*). Otherwise 0 is stored. Finally, the (*i, j, k*) yielding the maximum value in *PR*(*i, j, k*) is searched, and the corresponding *R_deleted_*(*i, j, k*) and *PR*(*i, j, k*) are obtained. The minimum PR from FVA for *R_deleted_*(*i, j, k*) are also calculated.

## 5.3 List of abbreviation

(FBA): Flux Balance Analysis
(FVA): Flux Variability Analysis
(LP): Linear Programming
(GR): Growth Rate
(PR): Production Rate
(TMGR): Theoretical Maximum Growth Rate
(TMPR): Theoretical Maximum Production Rate
(TMTF): Theoretical Maximum Total Flux
(GUR): Glucose Uptake Rate

## Declaration

### Ethics approval and consent to participate

Not applicable.

### Consent for publication

Not applicable.

### Availability of data and materials

All source codes and the solutions in the computational experiments described in this manuscript are freely available.

### Authors’ contribution

This work was done only by TT.

## Acknowledgements

The author appreciates the editor and reviewers for their efforts and valuable suggestions.

