## Supplemental file 1 for "Efficient reaction deletion algorithms for redesign of constraint-based metabolic networks for metabolite production with weak coupling": BTconstraintSearch.m

```
function [M,Brange,Trange,Prange,B,TMY,MB,rList,extype,targetRID,model2]=...
    BTconstraintSearch(model,targetMet,GUR,OUR,minGrowth,glucoseRxnID,oxygenRxnID,biomassRxnID,P)
%BTconstraintSearch obtains the constraints for the target production and
%biomass objective function, and returns the value of achieved target
%production.
%
%INPUTS
% model    COBRA model structure
% targetMet   target metabolite 
% GUR     glucose uptake ratio
% OUR     oxygen uptake ratio
% minGrowth    The minimum value of biomass objective fucnction
% glucoseRxnID     ID of glucose reaction in the COBRA model
% oxygenRxnID      ID of oxygen reaction in the COBRA model
% biomassRxnID     ID of biomass reaction in the COBRA model
% P           paramters for the cube size (described as $P^{-1}$ in the manuscript)
% 
%OUTPUTS
% M     achieved target metabolite production
% Brange     ID for the vertical axis of the cube that achieves the highest
%            target metabolite production
% Trange     ID for the horizontal axis of the cube that achieves the highest
%            target metabolite production
% B          The value of the biomass objective function when the highest
%            target metabolite production is achieved
% TMY         Theoretical Maximum yield (TMPR: Theoretical Maximum Production rate)
% MB          Maximum Biomass (TMGR: Theoretical Maximum Growth Rate)
%
%  Jan. 22, 2019   Takeyuki TAMURA

model.lb(glucoseRxnID)=-GUR;
model.lb(oxygenRxnID)=-OUR;
target=findMetIDs(model,targetMet);
m=size(model.mets,1);
n=size(model.rxns,1);
if isempty(find(strcmp(model.rxns,strcat('EX_',targetMet))))==0
    targetRID=find(strcmp(model.rxns,strcat('EX_',targetMet)));
    model2=model;
    extype=1;
elseif isempty(find(strcmp(model.rxns,strcat('DM_',targetMet))))==0
    targetRID=find(strcmp(model.rxns,strcat('DM_',targetMet)));
    model2=model;
    extype=2;
else
    [model2,rxnIDexists]=addReaction(model,'Transport',{targetMet},[-1]);
    m=size(model2.mets,1);
    n=size(model2.rxns,1);
    model2.S(target,n)=-1;
    model2.ub(n)=999999;
    model2.lb(n)=0;
    model2.rev(n)=0;
    targetRID=n;
    extype=3;
end

opt2=optimizeCbModel(model2);
MB=opt2.f;
model3=model2;
model3.c(biomassRxnID)=0;
model3.c(targetRID)=1;
model3.lb(biomassRxnID)=minGrowth;
opt2=optimizeCbModel(model3);
TMY=opt2.f;

lp.f=[zeros(n,1); -ones(n,1)];
lp.A=[-eye(n) -eye(n);eye(n) -eye(n)];
lp.b=[zeros(2*n,1)];
lp.Aeq=[model3.S zeros(m,n)];
lp.beq=zeros(m,1);
lp.lb=[model3.lb; zeros(n,1)];
lp.ub=[model3.ub; 999999*ones(n,1)];
options = optimoptions('linprog','Display','none');
[opt4.x, opt4.f, opt4.stat,opt4.output] = cplexlp(lp.f, lp.A, lp.b, lp.Aeq, lp.beq, lp.lb, lp.ub,options);
TMF=-opt4.f;


model3=model2;
model3.lb(biomassRxnID)=minGrowth;
lp.f=[zeros(n,1); ones(n,1)];
lp.A=[-eye(n) -eye(n);eye(n) -eye(n); zeros(1,n) -ones(1,n); zeros(1,n) ones(1,n)];
lp.Aeq=[model3.S zeros(m,n)];
lp.beq=zeros(m,1);
lp.lb=[model3.lb; zeros(n,1)];
lp.ub=[model3.ub; 999999*ones(n,1)];
tableBest=0;
biomassAtBest=0;
Brange=0;Trange=0;Prange=0;
rList={};
for i=1:P
    biomassLB=(MB/P)*(i-1);
    biomassUB=(MB/P)*i;
    for j=1:P
        targetLB=(TMY/P)*(j-1);
        targetUB=(TMY/P)*j;
        for k=1:P
        % for k=1:1
            %[i j k]
            tlb=(TMF/P)*(k-1);
            tub=(TMF/P)*k;
            %tlb=0;
            %tub=TMF;
            lp.b=[zeros(2*n,1); -tlb; tub];
            [table,opt4biomass,opt4target,opt4f,opt5biomass,rxnList]=...
                integrate(model2,model3,targetRID,biomassLB,biomassUB,targetLB,targetUB,n,biomassRxnID,lp);
            if (table > tableBest)&&(opt5biomass>=minGrowth)
                tableBest=table;
                biomassAtBest=opt5biomass;
                Brange=i;Trange=j;Prange=k;
                rList=rxnList;
            end
        end
    end
end
B=biomassAtBest;
M=tableBest;
%filename=sprintf('results/BTconstraintSearch_%d.mat',target);
%save(filename);
end
```
