## Supplemental file 1 for "Efficient reaction deletion algorithms for redesign of constraint-based metabolic networks for metabolite production with weak coupling": CubeProd.m

```
function [blockedRxns,  biomass,minFlux]...
    = CubeProd(varargin)
%CubeProd is a function of CubeProd that identifies a set of unused
%reactions for production of target metabolites.
%
%function [targetProduction, minFlux, maxFlux, blockedRxns, usedRxns, biomass]...
%    = CubeProd(model, targetMet,  glucoseRxn, oxygenRxn, biomassRxn, options)
%
%INPUTS
% model     COBRA model structure containing the following required fields to perform CubeProd.
%   rxns                    Rxns in the model
%   mets                    Metabolites in the model
%   S                       Stoichiometric matrix (sparse)
%   b                       RHS of Sv = b (usually zeros)
%   c                       Objective coefficients
%   lb                      Lower bounds for fluxes
%   ub                      Upper bounds for fluxes
%   rev                     Reversibility of fluxes
%
% targetMet   target metabolites
%             (e.g., {'succ_c','glu__D_c'} )
% glucoseRxn  Reaction representing glucose uptake (e.g., EX_glc__D_e)
% oxygenRxn   Reaction representing oxygen uptake (e.g., EX_o2_e)
% biomassRxn  Reaction representing biomass objective function
%                       (e.g., BIOMASS_Ec_iAF1260_core_59p81M)
% GUR    Glucose uptake ratio
% OUR    Oxygen uptake ratio
% minGrowth   The minimum value of biomass objective function that the
%             designed strain must achieve. 
% P           paramters for the cube size (described as $P^{-1}$ in the manuscript)
%
%OUTPUTS
% minFlux   The minimum values of the target metabolite production
%           obtained by FVA.
% blockedRxns   A set of unused reaction that achieves the value 
%               of targetProduction 
% biomass      The value of biomass objective function when blockedRxns
%              is not used.
%
%
% Jan. 22, 2019   Takeyuki TAMURA
%
s=size(varargin,2);
if size(varargin,2)<5
    error('''model'',''target'',''glucoseRxn'',''oxygenRxn''.''biomassRxn'' must be specified.')
end
model=varargin{1};
targetMet=varargin{2};
glucoseRxnID=findRxnIDs(model,varargin{3});
if glucoseRxnID==0
    error('invalid glucoseRxn name')
end
oxygenRxnID=findRxnIDs(model,varargin{4});
if oxygenRxnID==0
    error('invalid oxygenRxn name')
end
biomassRxnID=findRxnIDs(model,varargin{5});
if biomassRxnID==0
    error('invalid biomassRxn name')
end

for i=3:floor(s/2)
    if strcmp(varargin{2*i},'GUR')==1
        GUR=varargin{2*i+1};
    elseif strcmp(varargin{2*i},'OUR')==1
        OUR=varargin{2*i+1};
    elseif strcmp(varargin{2*i},'minGrowth')==1
        minGrowth=varargin{2*i+1};
    elseif strcmp(varargin{2*i},'P')==1
        P=varargin{2*i+1};
    else
        error('Options must be a subset of {type, GUR, OUR, minGrowth}')
    end
end


for i=1:1
       [minFlux(i),Brange(i),Trange(i),Prange(i),biomass(i),TMY(i),MB(i),blockedRxns,extype,targetRID,model2]=...
           BTconstraintSearch(model,targetMet{i},...
           GUR,OUR,minGrowth,glucoseRxnID,oxygenRxnID,biomassRxnID,P); 
end

save('CubeProd.mat');
end
```
