## Supplemental file 1 for "Efficient reaction deletion algorithms for redesign of constraint-based metabolic networks for metabolite production with weak coupling": example.m

```
function [outputArg1,outputArg2] = example(inputArg1,inputArg2)

load('iMM904.mat');
model=iMM904;
model.ub(359)=0;
model.lb(359)=0;
[blockedRxns,  biomass,minFlux]=...
CubeProd(model,{'urdglyc_c'},'EX_glc__D_e','EX_o2_e','BIOMASS_SC5_notrace','GUR',10,'OUR',2,...
'minGrowth',0.01,'P',5)

save('example.mat');
end
```
