## Supplemental file 1 for "Efficient reaction deletion algorithms for redesign of constraint-based metabolic networks for metabolite production with weak coupling": integrate.m

```
function [y,opt4biomass,opt4target,opt4f,opt5biomass,rxnList] =...
    integrate(model2,model3,targetRID,biomassLB,biomassUB,targetLB,targetUB,n,biomassRxnID,lp)
%'integrate' s a function of BTconstraintSearch.
%'integrate' employs linear programming twice in the designated cube.
%In the first LP, the unused reaction candidates are obtained.
%In the second LP, the production rate is calculated when reactions obtained in the first LP are not used.
%
%INPUTS
% model2   the model that a transport reaction is added to the original
%          model.
% model3   the model that the objective function has been modified in model2.
% targetRID  ID of the transport reaction of the target metabolite in
%            model2 and model3.
% biomassLB  the lower bound for the value of the biomass objective
%            function in the first LP.
% biomassUB  the upper bound for the value of the biomass objective
%            function in the first LP.
% targetLB   the lower bound for the value of the target metabolite
%            production in the first LP.
% targetUB   the upper bound for the value of the target metabolite
%            production in the first LP.
% n          the number of reactions
% biomassRxnID  ID of the reaction of biomass objectice function in model2
%               and model3
% lp         an linear programming formalization
%
%
%OUTPUTS
% y        the value of target metabolite production in the second LP. If
%          the LP is not feasible, -99999 is assigned.
% opt4biomass  the value of biomass objective function in the first LP.
% opt4target   the value of the target metabolite production in the first
%              LP.
% opt4f        the value of the objective function in the first LP.
%              Note that the biomass objective function is not optimized in
%              the first LP.
% opt5biomass  the value of biomass objective function in the second LP.
%
% Jan. 22, 2019   Takeyuki TAMURA
%

model4=model3;
lp.lb(biomassRxnID)=biomassLB;
lp.ub(biomassRxnID)=biomassUB;
lp.lb(targetRID)=targetLB;
lp.ub(targetRID)=targetUB;
%opt4=optimizeCbModel(model4);
options = optimoptions('linprog','Display','none','Algorithm','dual-simplex');
[opt4.x, opt4.f, opt4.stat,opt4.output] = cplexlp(lp.f, lp.A, lp.b, lp.Aeq, lp.beq, lp.lb, lp.ub,options);
if opt4.stat~=1
    y=-99999;
    opt4biomass=-99999;
    opt4target=-99999;
    opt4f=-99999;
    opt5biomass=-99999;
    rxnList={};
    return
end
opt4biomass=opt4.x(biomassRxnID);
opt4target=opt4.x(targetRID);
opt4f=opt4.f;
usedRxns=find(abs(opt4.x)>=0.0000001);
blockedRxns=setdiff(([1:n])',usedRxns);
rxnList=model4.rxns(blockedRxns);
model5=changeRxnBounds(model2,rxnList,0,'b');
opt5=optimizeCbModel(model5);
%[minFlux5,maxFlux5]=fluxVariability(model5,100,'max',{'Transport'});
[minFlux5,maxFlux5]=fluxVariability(model5,100,'max',model5.rxns(targetRID));
switch opt5.stat
    case 1
       % y=minFlux5.x(targetRID);
       y=minFlux5;
        opt5biomass=opt5.x(biomassRxnID);
    otherwise
        y=-999999;
        opt5biomass=-999999;
end
%display('end of the program')
return
end
```
